# Alpha Oscillatory Imbalance Predicts Attention Deficits in Visuospatial Neglect

**DOI:** 10.1101/2025.06.30.662471

**Authors:** Jelena Trajkovic, Marij Middag-van Spanje, Inge Leunissen, Teresa Schuhmann

**Affiliations:** Department of Cognitive Neuroscience, Faculty of Psychology and Neuroscience, Maastricht University, 6229 ER, The Netherlands; InteraktContour, 8070 AC Nunspeet, The Netherlands

**Author notes:** **Correspondance :** Jelena Trajkovic.

**Keywords:** Visuospatial neglect, EEG resting state, spatial attention, alpha oscillations

## Abstract

**Background and Objectives:** Visuospatial neglect (VSN) is a common consequence of unilateral stroke, characterized by reduced awareness on the contralesional side of space. This study examines resting-state electrophysiological mechanisms underlying VSN, with a focus on interhemispheric differences in alpha-band oscillations and their association with deficits in visual perception and attention.

**Methods:** Resting-state EEG was recorded during three-minute eyes-closed and eyes-open periods in VSN patients. Participants also completed a battery of perceptual and attentional assessments, including the Star Cancellation Task, the Computerized Visual Detection Task, and the Line Bisection Task. Measures of alpha activity, specifically amplitude, peak frequency, and fronto-parietal connectivity,were compared between VSN patients and age- and gender-matched healthy controls. Within the VSN group, alpha activity in the lesioned and non-lesioned hemispheres was used to predict task performance.

**Results:** Compared to healthy controls, VSN patients displayed a distinct resting-state alpha profile, characterized by slower and less synchronized alpha oscillations and reduced fronto-parietal alpha connectivity in the lesioned hemisphere. In contrast, controls showed symmetrical alpha activity across hemispheres. Crucially, the degree of interhemispheric alpha asymmetry in VSN patients consistently predicted attentional impairments across all behavioral tasks.

**Discussion:** VSN is linked to disrupted alpha oscillatory dynamics and interhemispheric imbalance. The extent of alpha desynchronization between hemispheres predicts symptom severity, supporting the interhemispheric competition model of attention. These findings suggest that hyperactive alpha activity in the non-lesioned hemisphere may suppress the lesioned hemisphere, contributing to attentional bias away from the contralesional space.

## Introduction

Visuospatial neglect (VSN) is a prevalent neurocognitive disorder characterized by a patient’s inability to attend to and report stimuli appearing in the contralesional hemispace ^1–3^. This condition most commonly arises following a stroke affecting cortical and subcortical brain regions, particularly within the right frontoparietal network, which plays a key role in spatial attention. Although neglect can result from lesions in either hemisphere, damage to the right hemisphere is more often associated with severe and persistent attentional deficits ^4–7^. This asymmetry has contributed to the widely accepted hypothesis of right hemisphere dominance in spatial attention ^8,9^. This theory presumes that the left hemisphere is responsible for directing attention toward the right side of space, whereas the right hemisphere governs attention shifts toward both the left and right sides ^10^. As a result, damage to the right hemisphere leaves only the left hemisphere’s limited capacity to attend to the right visual field, leading to pronounced neglect of the left side. In contrast, lesions in the left hemisphere tend to cause milder neglect, as the intact right hemisphere retains the ability to direct attention across both visual fields.

In the healthy brain, visual attention relies on coordinated activity between “control” regions in the frontal and parietal cortices, and sensory regions in the occipital cortex. These interactions enhance perception by guiding attentional focus and generating expectations about visual input ^2,11,12^. A possible physiological marker of the functional interaction between frontoparietal regions and occipital visual areas is the modulation of posterior alpha rhythms, measured using electroencephalography (EEG). In the absence of visual input, parieto-occipital regions typically exhibit strong alpha synchronization. However, in preparation for visual targets, these rhythms dynamically desynchronize ^13–16^.

During rest, posterior alpha activity reflects baseline cortical excitability, and imbalances in this excitability across hemispheres may contribute to the attentional biases observed in VSN. Specifically, right-hemisphere lesions may lead to a dual disruption: decreased alpha activity in the damaged hemisphere and disinhibition of homologous areas in the intact left hemisphere ^4,9,17^. This imbalance, marked by increased left-hemisphere alpha activity, may further suppress the already impaired right hemisphere ^6,13,18^ , effectively shifting attention toward the ipsilesional (right) space.

Recent studies have shown that not only alpha amplitude but also the speed of alpha oscillations, defined as the frequency peak within the alpha band, is crucial in the perceptual processing of sensory stimuli. Specifically, faster alpha oscillations result in higher perceptual sampling rates and more accurate perceptual experiences^15,19–25^. Conversely, alpha peak frequency is often reduced after stroke, which has been associated with various perceptual and cognitive abnormalities, possibly contributing to attention deficits in VSN.

Finally, effective attention shifting depends on interregional alpha phase connectivity, particularly between prefrontal and posterior contralateral areas^26^. This suggests that the prefrontal cortex may coordinate posterior alpha dynamics in a top-down fashion. Supporting this, resting-state MRI studies in VSN patients show decreased fronto-parietal connectivity in the lesioned hemisphere^27^, while EEG studies highlight that reduced functional connectivity, both at rest^28^ and during attention tasks^29^, is specific to the alpha frequency range and correlates with impaired attentional performance. Thus, unilateral lesions affecting frontal or parietal cortices may impair the flexible modulation of the alpha system, thereby contributing to spatial attentional deficits in VSN.

The present study aims to characterize the resting-state electrophysiological profile of patients with visuospatial neglect. Understanding this baseline activity is particularly important given the high prevalence and clinical impact of VSN, which is often underdiagnosed due to variability in its presentation and the limitations of current assessment tools^30^. Traditional paper-and-pencil assessments remain widely used but may lack sensitivity. Severely affected patients may be unable to complete such tasks, while mildly affected individuals may score within the normal range^31^. This underscores the need for complementary assessment methods that can detect subtle physiological changes, such as EEG-based biomarkers.

Based on previous findings in both healthy ^15,26,32,33^ and neglect patients ^29^, the expected EEG profile of such patients includes a lower alpha amplitude, slower alpha frequency and reduced fronto-parietal connectivity in the lesioned hemisphere. Our decision to focus on resting-state EEG, rather than task-based recordings, is motivated by both theoretical and practical factors. Theoretically, since VSN emerges after stroke, pathological changes should already be apparent during rest, offering a window into the altered baseline dynamics of the brain. Practically, resting-state EEG is a simple and versatile method, making it an effective screening tool that allows for task-independent, standardized measurements in large-scale assessments.

If we can identify specific resting-state EEG signatures that reliably predict the severity of attentional deficits, these biomarkers could be leveraged to enhance VSN diagnosis, quantify symptom severity, and potentially guide prognosis and treatment planning.

## Methods

### Participants

Twenty-two chronic stroke patients (6 F, Age: M = 60.09 years, all with right-hemispheric lesion, sample size estimate available at ^34^) were recruited for the current study from healthcare organizations in The Netherlands that are specialized in supporting and treating people with brain injury. Patient recruitment was performed by a psychologist.

Inclusion criteria were: 1. Neurologically objectified stroke 2. Stroke occurring at 18-80 years of age 3. Stroke occurring at least six months ago 4. Sufficient capacity of communication and comprehension (as assessed by a psychologist) 5. Presence of VSN confirmed with a screening. Exclusion criteria included: 1.

Current participation in alternative neglect treatment as to avoid potential cross-contamination (as the participants were included enrolled in the Tacs clinical trial) 2. Physical or mental inability to participate (as assessed by a psychologist) 3. Presence of hemianopia (based on clinical judgment) 4. Presence of a diagnosed (neuro)psychiatric or neurodegenerative disease 5. Substance abuse 6. Pregnancy. ^35^To compare the EEG activity of the stroke patients with a normative EEG profile, we also included 22 age-(M_VSN_= 60.09; M_HC_= 60.04; t(42) = 0.059, p = .953) and gender-matched (χ^2^ = 0.0; *p = 1.00*) healthy controls, pseudo-randomly selected from the online repository (doi:10.18112/openneuro.ds005385.v1.0.0). These EEG measurements are part of the Dortmund Vital Study, an ongoing longitudinal cohort study on the development of cognitive functions over an age range from 20 to 70 years, conducted by the Leibniz Research Centre for Working Environment and Human Factors at the Technical University Dortmund (IfADo) (ClinicalTrials.gov Identifier: NCT05155397).

**Table 1.** Descriptive Statistics. VSN: Visuo-spatial Neglect ; HC: Healthy Controls; F: Female.

|  | Age |  | Gender |  |
| --- | --- | --- | --- | --- |
|  | VSN | HC | VSN | HC |
| Mean | 60.091 | 60.045 | 27% F | 27%F |
| Std. Deviation | 11.002 | 9.791 |  |  |

### Standard Protocol Approvals, Registrations, and Patient Consents

The study was approved by the Medical-Ethical Committee azM/UM of Maastricht University (NL70256.068.19 / METC 19-047). All participants gave written informed consent before taking part in the study. The here-reported data is part of the clinical trial registered in <u>ClinicalTrials.gov</u> (Non-Invasive Brain Stimulation as an Innovative Treatment for Chronic Neglect Patients (NibsNeglect), NCT05466487).

### Experimental Procedure

For the VSN, EEG resting-state measurements were obtained during 3-minute resting state for eyes-closed and eyes-opened conditions. Behavioral measures during the computerized visual detection task (CVDT) were also obtained, together with the Star Cancellation task (SCT) and Schenkenberg line bisection tasks (SLBT).

Data for the HC also consists of the 3 minutes eyes-opened and eyes-closed resting-state measurements^36^.

### Behavioral outcome: Computerized visual detection task (CVDT)

The CVDT (Figure 1A) evaluates perceptual sensitivity and attentional selection in each hemifield individually, as well as in the context of competing visual stimuli across both hemifields. It has been shown to be a reliable tool for assessing unilateral neglect and extinction ^35,37,38^. During the task, patients were seated in front of a computer screen. They were asked to fixate on the center of the screen, marked with a fixation cross. Gabor patches were displayed on the left, right, and both sides of the screen (40 trials per location). The patient had to identify the stimulus location by pressing the <, >, or ˅ key accordingly. The contrast of the stimuli was adjusted adaptively on a trial-by-trial basis for each of the three locations independently using the QUEST staircase algorithm ^39^. In the offline analysis, we calculated and averaged the hit rate for the contralesional stimuli and bilateral stimuli. In order to account for the fact that contrast levels varied on a trial-by-trial basis, thus factoring in task difficulty, hits were weighted by the contrast level, according to the following formula: **x = log10(max_contrast) / log10(trial_contrast)**. This results in a potential scoring range of 0 to 76.49 weighted hits per condition.

**Figure 1.**
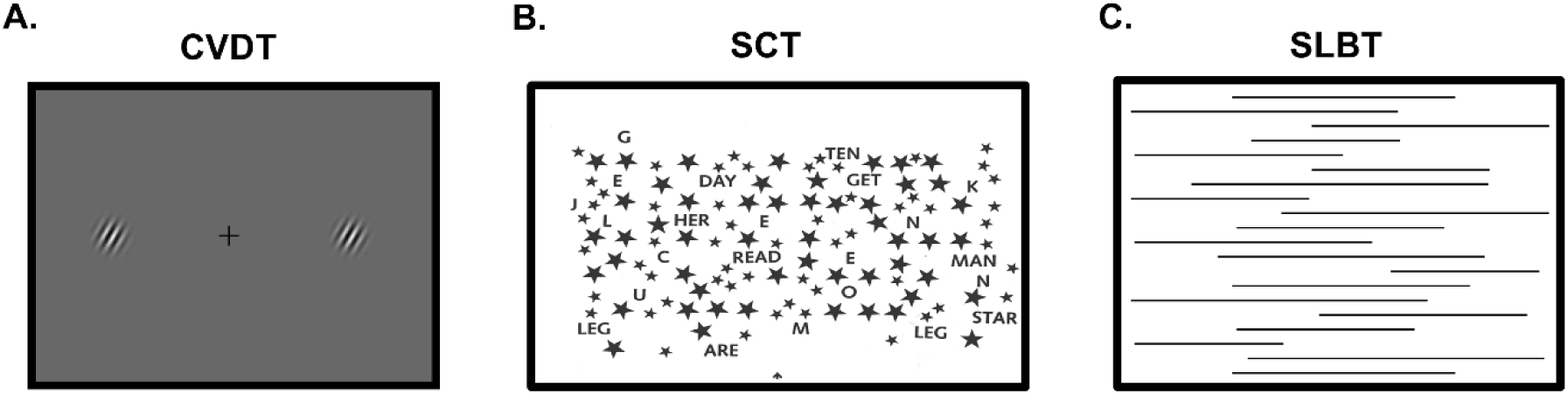
Behavioral Outcomes. **A.** Computerized visual detection task (CVDT). Participants were instructed to detect the Gabor Patches appearing on the left, right or bilaterally (as depicted in the figure), while fixating the center of the screen. **B.** Star Cancellation Task (SCT). Participants were instructed to identify and mark all target items (small stars) by touching the screen with their finger. **C.** Schenkenberg line bisection task (SLBT). Participants were instructed to indicate the perceived midpoint of each line.

### Behavioral outcome: Star cancellation task (SCT)

The SCT (Figure 1B) included 52 large stars, 13 letters, and 10 short words, interspersed with 56 smaller stars^40^, all displayed on a laptop screen. The patient was instructed to identify and mark all target items (small stars) by touching the screen with their finger. The Quality of Search (QoS) score integrates both accuracy and speed into a single metric, representing the optimal accuracy-to-speed search ratio, calculated using the following formula ^41^:

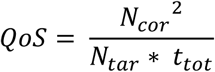

where QoS is the quality of search score, N_cor_ is the number of cancelled targets, N_tar_ is the total number of targets, and t_tot_ is the total time spent.

A higher score indicates a greater number of correctly marked targets combined with a faster cancellation speed ^41^.

### Behavioral outcome: Schenkenberg line bisection task (SLBT)

The SLBT (Figure 1C) comprised 20 horizontal lines, ranging from 10 to 20 cm in length, positioned at three different locations (left, middle, and right) on a landscape-oriented A4 sheet. The patient was instructed to indicate the perceived midpoint of each line. Relative deviation scores were then calculated and averaged per line position to generate the left, middle, and right average scores:

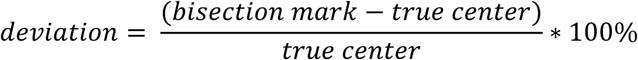

where values are always measured from the left end of the line.

The analysis focused solely on lines located on the contralesional side, where the poorest performance was expected.

### EEG recording

A set of 32 Ag/AgCl electrodes was mounted according to the international 10–20 system (BrainCap). Additionally, four EoG channels were positioned: on the outer canthi of both eyes, as well as above and below the left eye. The right and left mastoids were used as the online reference. Ground was positioned on the right cheek of the subject. All impedances were kept below 10 kΩ. EEG signals were recorded with a pass band filter 0.5–250 Hz at a sampling rate of 1000 Hz.

For the HC, the EEG was measured using a 64-channel elastic cap arranged based on the 10–20 system with FCz electrode as on-line reference. The EEG signal was recorded with a 1000-Hz sampling rate and filtered online by a 250-Hz low-pass filter. No online high-pass filter was used. Impedances were kept below 10 kΩ ^36^.

All EEG analyses were implemented by custom-made routines developed in Matlab R2022b (The MathWorks, Inc., Natick, Massachusetts, United States) using EEGLab toolbox functions (v. 2023.0).

### EEG analysis

Processing of the eyes-open and eyes-closed resting-state data for both groups included re-referencing of the signal to the average activity across all electrodes and an offline resampling to 500 Hz. After that, the data was split in 1 second epochs. Four EoG electrodes (for VSN patients) and frontal electrodes (for HC) were used to automatically detect and remove the epochs with outlier activity containing eye movements. Finally, data were visually inspected to remove the remaining noisy trials.

*Alpha amplitude* values were determined by spectral analysis. Specifically, complex Morlet wavelet convolution was applied to the signal, with Morlet wavelet peak frequencies ranging from 5 to 65Hz in 60 logarithmic steps. The full width at half-maximum (FWHM) ranged from 500 to 200ms with increasing wavelet peak frequency. Data was then averaged for the alpha frequency range (7-13 Hz), over posterior electrodes, and separately for the left (CP1, CP5, P3, P7 and O1) and right sensors (CP2, CP6, P4, P8 and O2).

*Individual alpha frequency* was calculated by first calculating power spectrum density using Welch’s method, with a Hanning windows tapering with 10% overlap, and the resulting 0.1 frequency resolution. The spectra were then decomposed into periodic and aperiodic components using the FOOOF algorithm. The obtained flattened spectra (periodic-aperiodic component) in the alpha range (7-13 Hz) was fitted to a Gaussian curve using Matlab fit function. The Gaussian model fits peaks, and is given by:

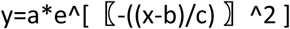

where b is the centroid, which we used as an estimation of the IAF ^42^. IAF was computed separately for the left (CP1, CP5, P3, P7 and O1) and right sensors (CP2, CP6, P4, P8 and O2).

*Alpha connectivity* was estimated in the sensor space between frontal and posterior electrodes of both hemispheres (frontal electrodes: FC2/FC1, FC4/FC5, FP2/FP1, F8/F7, F4/F3; posterior electrodes: CP2/CP1, CP4/CP5, P4/P5, P8/P7, O2/O1) via the weighted phase-lag index (wPLI). This is a measure of phase lag-based connectivity, which accounts for non-zero phase lag/lead relations between two signal time series, as it defines connectivity as the absolute value of the average sign of phase angle differences and additionally de-weighting vectors that are closer to the real axis such that those vectors have a smaller influence on the final connectivity estimate. By extension, this measure is insensitive to volume conduction and noise of a different origin, considered optimal for exploratory analysis as it minimizes type-I errors ^43^. To obtain wPLI values, time series data was first transformed into the time-frequency domain via convolution with a family of complex Morlet wavelets (the number of cycles increased from 5 to 18 in logarithmic steps). Therefore, for frequencies ranging from 3 to 30Hz in 1Hz steps, first convolution by frequency-domain multiplication was performed and then the inverse Fourier transformation was taken. Phase was defined as the angle relative to the positive real axis, and phase differences were then computed between the chosen electrode pairs. Finally, wPLI was calculated as the absolute value of the average sign of phase angle differences, whereby vectors closer to the real axis were de-weighted. Finally, wPLI values were averaged for alpha frequency band (7-13 Hz).

### Statistical analysis

To compare resting-state EEG activity between the VSN and healthy control (HC) groups, we conducted a series of 2×2 mixed-design ANOVAs for each alpha parameter: alpha amplitude, individual alpha frequency, and fronto-posterior alpha connectivity. Each analysis included a between-subjects factor of GROUP (VSN, HC) and a within-subjects factor of HEMISPHERE (LEFT, RIGHT). Planned follow-up comparisons between hemispheres within each group were conducted using two-tailed Student’s t-tests.

To assess whether the alpha activity profile could predict attentional deficits in VSN patients, we used a backward linear regression analysis. In this model, task performance was the dependent variable, and all alpha parameters from both hemispheres served as predictors. The analysis began with a full model including all predictors, then iteratively removed the least significant variables to arrive at the most parsimonious model that retained strong explanatory power.

### Data availability

The data that support the findings of this study are available on request from the corresponding author.

## Results

### Interhemispheric differences in alpha activity in neglect patients

To examine the interhemispheric differences in alpha oscillatory parameters, we analyzed alpha amplitude and frequency in the parietal cortex and fronto-parietal connectivity of neglect patients (N=22) and the age- and gender-matched healthy control group (N=22). Specifically, we compared the interhemispheric differences (RIGHT, LEFT HEMISPHERE) in the alpha activity across the two groups (HC, VSN) via 2-way mixed ANOVA design.

#### Alpha amplitude

During the eyes-closed condition, we observed a significant GROUP × HEMISPHERE interaction for alpha amplitude (F(1,42) = 4.727, *p = .035*, ηp² = 0.101, Figure 2A), indicating that the interhemispheric distribution of alpha amplitude differed between groups. In HC, alpha amplitude was symmetrically distributed across hemispheres (M_right_ = 1.973, M_left_ = 1.991; t(21) = 0.386, *p = .703*, d = 0.15, Figure 4A). In contrast, VSN patients showed marked asymmetry, with significantly higher alpha synchronization in the left (non-lesioned) hemisphere (M_right_ = 2.031, M_left_ = 3.368; t(21) = 2.141, *p = .044*, d = 0.46, Figure 3A). No significant group or interhemispheric differences in alpha amplitude were found during the eyes-open condition (all F_s_ < 2.034, all *p_s_ > .161*, all ηp²_s_ < 0.046, Figure 3A and Figure 4A).

**Figure 2.**
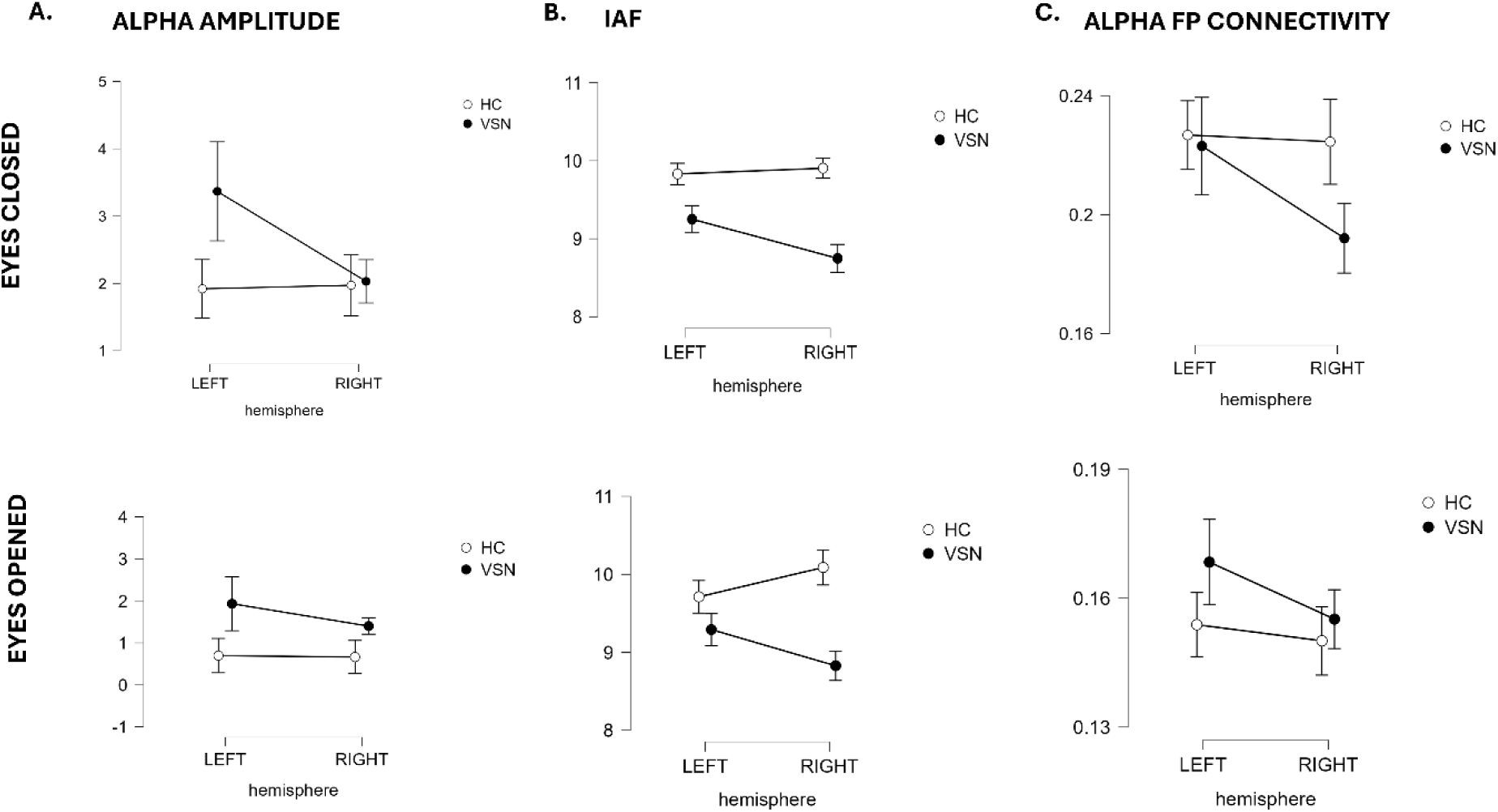
Interhemispheric differences in alpha parameters of VSN patients (lack dots) and healthy controls (HC; white dots) during eyes-closed (top panel) and eyes-open rs-EEG (bottom panel). **A.** Alpha amplitude differences between the HC and VSN patients across the two hemispheres. **B.** Individual Alpha Frequency (IAF) differences between the HC and VSN patients across the two hemispheres. **C.** Fronto-posterior (FP) alpha connectivity differences between the HC and VSN patients across the two hemispheres. Group means are represented by white and black dots, while error bars represent standard error of the mean.

**Figure 3.**
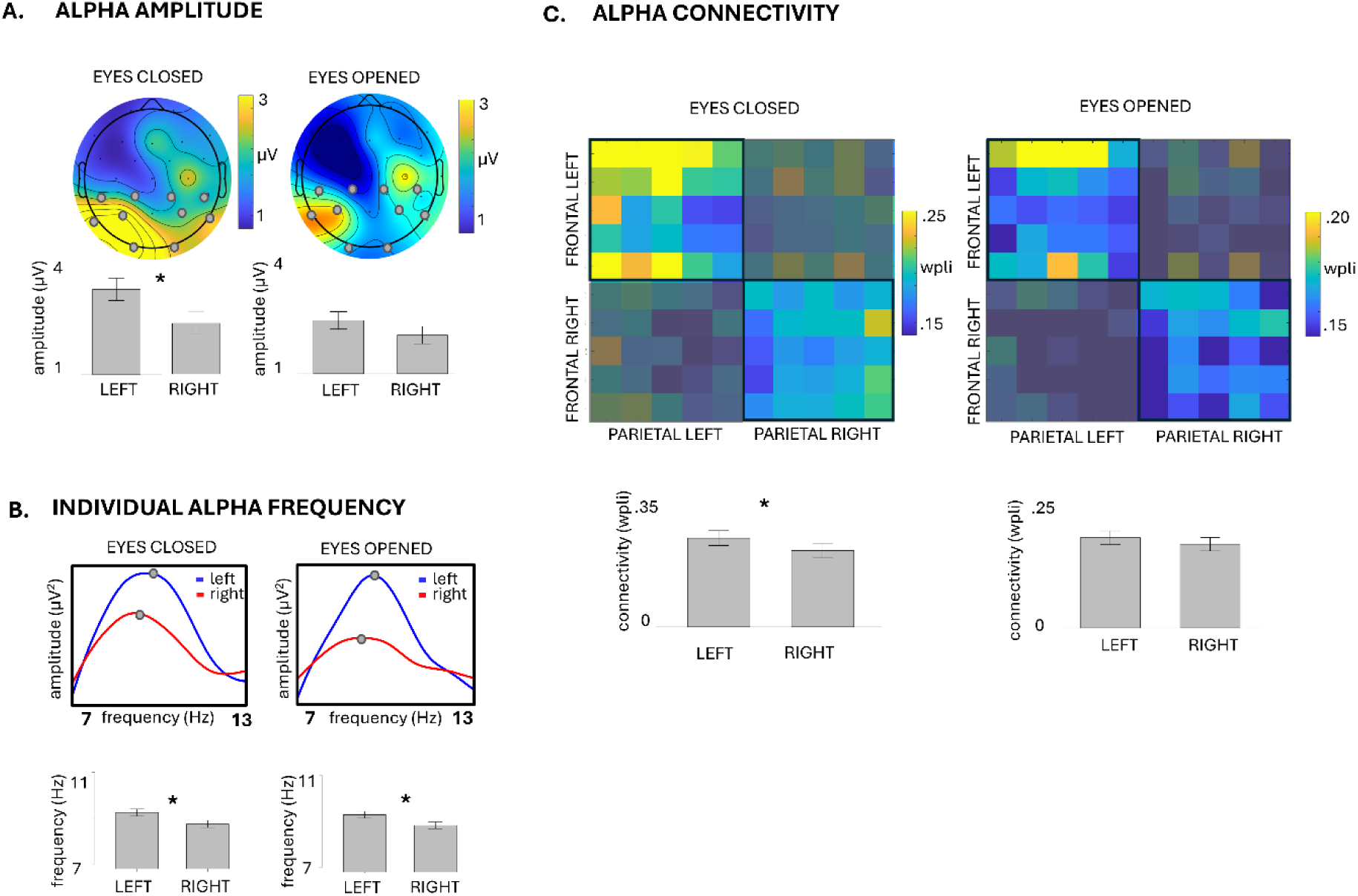
VSN patients: Interhemispheric differences in alpha activity. **A.** Alpha amplitude: Top: Topographies of the scalp distribution of the average alpha amplitude (7-13 Hz) during the eyes-closed and eyes-opened resting state. Grey circles mark the electrodes that were used for the analysis. Bottom: Bar plots of the alpha-amplitude levels in the left and right parietal sensors, during eyes-closed and eyes-opened resting state condition. **B.** Individual Alpha Frequency (IAF): Top: Average power spectrum densities (PSD) of the left (in blue) and right (in red) ROI (Region of Interest) for the eyes-closed and eyes opened resting state condition. Grey circles indicate the IAF. Bottom: Bar plots of the alpha-frequency in the left and right parietal sensors, during eyes-closed and eyes-opened resting state condition. **C.** Alpha Connectivity: Top: Matrices of the weighted Phase Lag Index (wPLI) for the frontal and parietal regions of the right and left hemisphere for the eyes-closed and eyes opened resting state condition. Bottom: Bar plots of the fronto-parietal connectivity strength in the left and right frontal and parietal sensors, during eyes-closed and eyes-opened resting state condition. Two-tailed t-test statistical significance is reported ( ∗ p *<* 0.05). Error bars represent standard error of the mean, **μ** v = microvolt, Hz = Hertz.

**Figure 4.**
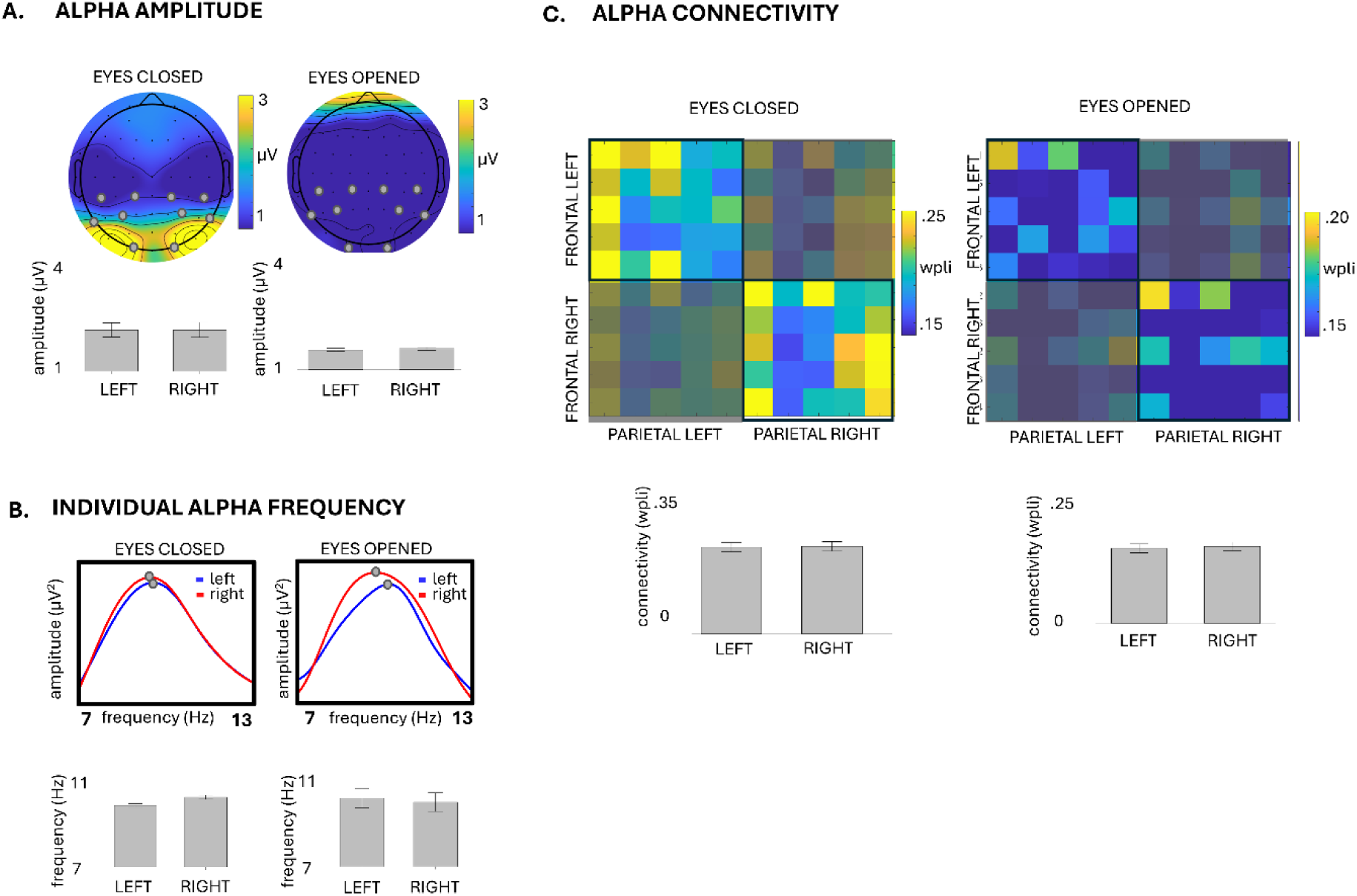
Healthy Controls: Interhemispheric differences in alpha activity. **A.** Alpha amplitude: Top: Topographies of the scalp distribution of the average alpha amplitude (7-13 Hz) during the eyes-closed and eyes-opened resting state. Grey circles mark the electrodes that were used for the analysis. Bottom: Bar plots of the alpha-amplitude levels in the left and right parietal sensors, during eyes-closed and eyes-opened resting state condition. **B.** Individual Alpha Frequency (IAF): Top: Average power spectrum densities (PSD) of the left (in blue) and right (in red) ROI (Region of Interest) for the eyes-closed and eyes opened resting state condition. Grey circles indicate the IAF. Bottom: Bar plots of the alpha-frequency in the left and right parietal sensors, during eyes-closed and eyes-opened resting state condition. **C.** Alpha Connectivity: Top: Matrices of the weighted Phase Lag Index (wPLI) for the frontal and parietal regions of the right and left hemisphere for the eyes-closed and eyes opened resting state condition. Bottom: Bar plots of the fronto-parietal connectivity strength in the left (P3 and F3) and right frontal and parietal sensors, during eyes-closed and eyes-opened resting state condition. Two-tailed t-test statistical significance is reported ( ∗ p *<* 0.05). Error bars represent standard error of the mean, **μ** v = microvolt, Hz = Hertz.

#### Alpha frequency

We found that IAF differences between groups were hemisphere-specific, as indicated by a significant GROUP × HEMISPHERE interaction in both the eyes-closed (F(1,42) = 5.843, *p = .020*, ηp² = 0.122, Figure 2B) and eyes-open conditions (F(1,42) = 6.203, *p = .017*, ηp² = 0.129, Figure 2B). In healthy controls, alpha frequency was similar across hemispheres (eyes closed: M_right_ = 9.904, M_left_ = 9.830; t(21) = 0.807, *p = .429*, d = 0.18; eyes open: M_right_ = 10.088, M_left_ = 9.711; t(21) = 1.439, *p = .165*, d = 0.31, Figure 4B). In contrast, VSN patients exhibited significantly faster alpha oscillations in the non-lesioned (left) hemisphere compared to the lesioned (right) hemisphere in both conditions (eyes closed: M_right_ = 8.748, M_left_ = 9.249; t(21) = 2.277, *p = .033*, d = 0.48; eyes open: M_right_ = 8.827, M_left_ = 9.291; t(21) = 2.182, *p = .041*, d = 0.46, Figure 3B).

#### Fronto-parietal connectivity

Finally, fronto-parietal connectivity during the eyes-closed condition showed a marginally significant GROUP × HEMISPHERE interaction (F(1,42) = 3.445, *p = .070*, ηp² = 0.076, Figure 2C), indicating group-specific hemispheric differences. In healthy controls, alpha connectivity was balanced across hemispheres (M_right_ = 0.225, M_left_ = 0.227; t(21) = 0.273, *p = .787*, d = 0.17, Figure 4C). In contrast, VSN patients demonstrated reduced alpha connectivity in the lesioned (right) hemisphere compared to the non-lesioned side (M_right_ = 0.191, M_left_ = 0.223; t(21) = 2.367, *p = .028*, d = 0.38, Figure 3C). These effects were specific to the eyes-closed condition, with no significant group or hemispheric differences observed during eyes-open recording (F(1,42) = 0.819, *p = .371*, ηp² = 0.019, Figure 3C and Figure 4C).

In sum, resting-state profile of the alpha oscillatory activity is different between the VSN patients and age- and gender-matched healthy controls. While the alpha amplitude, alpha frequency and fronto-posterior connectivity in the alpha range are comparable between the hemispheres in healthy participants, VSN displayed a higher alpha amplitude, faster alpha frequency and higher connectivity in the non-lesioned hemisphere compared to the lesioned one. These differences were particularly evident during eyes-closed resting state condition, and less visible during eyes-opened condition.

#### Exploration of adjacent frequency bands (Theta and beta)

For completeness, we also investigated whether interhemispheric differences in VSN patients are also present in other adjacent frequency bands, mainly theta and beta frequency. While in the theta band we did not find any differences in connectivity values or amplitude between the two hemispheres (all ts<2.021, all *p*s < .06), in the beta band there was again a significant trend of higher amplitude in the left compared to the right hemisphere during both eyes-open (M_right_= 0.400; M_left_= 0.621; t(21) = 2.375, p = .027; d = 0.54) and eyes-closed (M_right_= 0.288; M_left_= 0.369; t(21) = 2.517, p = .020; d = 0.51) condition, without connectivity differences (all ts<1.765, all *p*s < .09).

#### Resting-state alpha activity predicts attentional deficits

Next, we investigated whether resting-state alpha parameters could predict attentional deficits in neglect patients across different tasks.

### Computerized Visual Detection Task (CVDT)

While the eyes-open resting state parameters could not predict behavioral performance, a model based on the eyes-closed alpha-amplitude parameters predicted the attentional bias (R^2^ = 0.491, F (21) = 5.778, p = .006). Specifically, a higher alpha-amplitude in the right hemisphere (β = 1.335, t = 3.319, p = .004), along with the lower alpha amplitude in the left hemisphere (β = -1.298, t = 3.226, p = .004), predicted a higher perceptual accuracy for the stimuli appearing in the neglect hemifield, or bilaterally (Figure 5).

**Figure 5.**
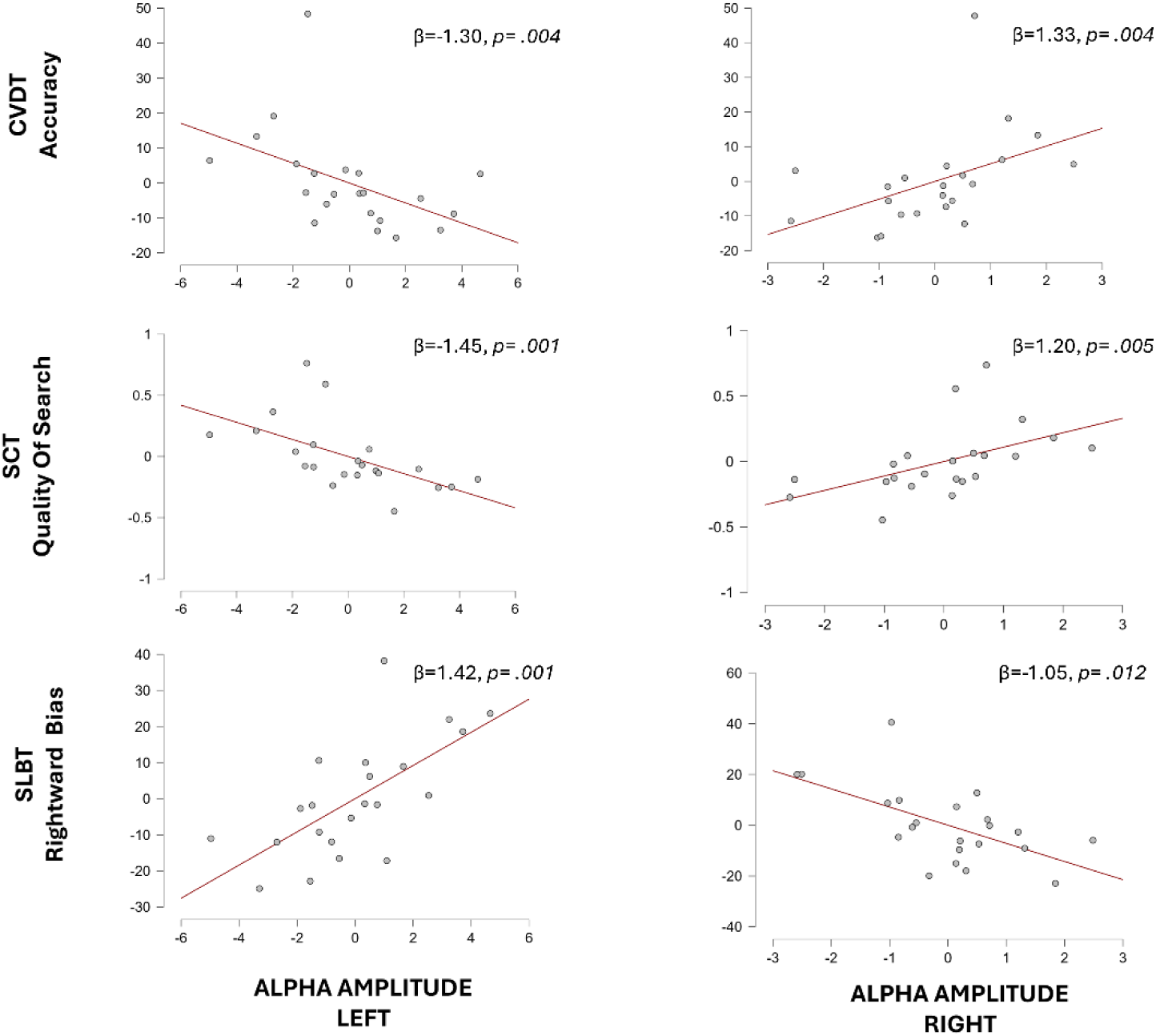
Alpha amplitude and task performance on attention tasks. Results of the regression model predicting the performance during the Computerized Visual Detection Task (CVDT, top panels) for the left and bilateral stimuli, Stat Cancellation Task (SCT, middle panels) and the Schenkenberg line bisection task (SLBT, bottom panels) based on the alpha amplitude levels in the right (right panels) and left hemisphere (left panels).

Additionally, higher connectivity in the right (lesioned) hemisphere was a marginally significant predictor of a higher performance on the same task (β = 0.344, t = 2.028, p = .058). Other predictors, such as individual alpha frequency and the connectivity in the left hemisphere were not predictive of task performance.

### Start cancellation task

Similar results were also obtained for the star cancellation task, where, again, interhemispheric differences in the alpha-amplitude predicted the quality of search score (R^2^ = 0.437, F (21) = 7.379, p = .004). Specifically, a higher alpha synchronization in the lesioned hemisphere (β = 1.203, t = 3.153, p = .005) and lower alpha amplitude in the non-lesioned one (β = -1.451, t = 3.804, p = .001) during eyes-closed condition predicted better task performance on the SCT (Figure 5). Along with the alpha-amplitude distribution, IAF measured during eyes-open condition was also a significant predictor of the quality of search score (R^2^ = 0.546, F (21) = 5.116, p = .007), with faster alpha in the right hemisphere (β = 0.950, t = 3.725, p = .002) and slower alpha in the left (β = -0.728, t = 2.726, p = .014) predicting better performance. On the other hand, resting-state alpha connectivity was not a significant predictor of attention efficiency.

### Line bisection task

Finally, for the line bisection task, the deviation in bisection of the lines presented in the left hemifield, as measured with the Schenkenberg line bisection task (SLBT) was exclusively predicted by the asymmetry in the levels of alpha-amplitude across the two hemispheres in both eyes-closed and eyes-open resting state condition (eyes-closed: R^2^ = 0.451, F (21) = 7.818, p = .003; eyes-opened: R^2^ = 0.311, F (21) = 4.294, p = .029), with a lower alpha in the left (non-lesioned) (eyes-closed: β = 1.415, t = 3.757, p = .001; eyes-opened: β = 2.171, t = 2.365, p = .029) along with a higher alpha in the right (lesioned) (β = - 1.053, t = 2.797, p = .012; eyes-opened: β = -1.794, t = 1.954, p = .066) hemisphere predicting less rightward bias during line bisection (Figure 5).

### Specificity of alpha effects compared to theta and beta oscillations

Crucially, this pattern of significant predictors across various task contexts was consistent only for the alpha band, but not the theta or beta frequency. Specifically, while in theta no significant predictors were found across any of the tasks (all R^2^s < 0.169, all Fs (21) < 4.097, all ps > .057), a higher beta connectivity, along with a lower beta amplitude in the left hemisphere during eyes-open (but not eyes-closed, all R^2^s < 0.158, all Fs (21) < 3.743, all ps > .067) condition predicted better task performance on star cancellation task (R^2^ = 0.536, F (21) = 6.927, p = .003) and the CVDT task (R^2^ = 0.402, F (21) = 6.379, p = .008).

## Discussion

Understanding the electrophysiological basis of attentional deficits in patients with visuospatial neglect (VSN) is essential not only for deepening our insight into the oscillatory mechanisms of spatial attention but also for guiding potential treatment strategies. To this end, we recorded resting-state EEG in patients with VSN and in age- and gender-matched healthy controls (HC). Neglect severity was assessed using the Computerized Visual Detection Task (CVDT), Star Cancellation Task (SCT), and Line Bisection Task. Our main objective was to examine interhemispheric differences in alpha activity during both eyes-open and eyes-closed resting states and to explore how these neural markers relate to attentional biases observed across behavioral tasks.

### Interhemispheric imbalance in alpha activity as a marker of attentional bias

Our results revealed distinct patterns of alpha amplitude distribution between VSN patients and healthy controls. As expected, healthy participants showed a symmetrical distribution of resting-state alpha amplitude across both hemispheres, consistent with previous findings ^29,44–49^. In contrast, VSN patients displayed significant interhemispheric differences during the eyes-closed condition, characterized by higher alpha synchronization in the non-lesioned (left) hemisphere. This pattern replicates prior reports of alpha asymmetries in neglect^29^. Crucially, we found that this asymmetry predicted attentional bias across all administered tasks: greater alpha amplitude in the left hemisphere, coupled with reduced amplitude in the lesioned right hemisphere, was associated with stronger attentional bias toward the ipsilesional (right) space. These findings support theoretical models suggesting that neglect stems from interhemispheric imbalance in cortical excitability ^4,17,50^. Specifically, right hemisphere lesions not only suppress activity within the damaged network but also lead to disinhibition and compensatory hyperactivity in the left hemisphere ^9^. This compensatory overactivation may exacerbate imbalance by further inhibiting the already weakened right hemisphere ^6,14,18,29,50,51^. Our results reinforce this framework, highlighting that interhemispheric alpha asymmetries at rest are a robust neural correlate of spatial attention deficits in VSN.

### Slower alpha frequency and reduced fronto-parietal connectivity in the lesioned hemisphere

In addition to alpha amplitude asymmetries, we also found that VSN patients exhibited a slower alpha frequency and reduced fronto-parietal connectivity in the lesioned hemisphere—patterns not observed in healthy controls, who showed a symmetrical distribution of these alpha parameters across hemispheres. These findings are consistent with prior research. Specifically, a recent study showed that patients with left visuospatial neglect have a slower alpha frequency and lower fronto-parietal connectivity in the right hemisphere during the task baseline (i.e., fixation cross) ^29^. Slower IAF has been linked to a lower ability to accurately perceive sensory information and less efficient sensory sampling ^15,19–21^. In the context of visual spatial attention, efficiently focusing attention to one hemispace has been associated with a faster contralateral IAF, and slower IAF in the ipsilateral hemisphere ^15^. Therefore, slower alpha-activity in the lesioned hemisphere of the neglect patients during rest could indicate their attentional bias toward ipsilateral hemispace.

Similarly, a higher fronto-parietal connectivity has been shown during attentional demands ^52,53^, predicting higher perceptual sensitivity and metacognitive abilities ^54,55^. Consequently, lower connectivity in the lesioned hemisphere of the neglect patients could relate to an impaired processing of contralesional stimuli. In sum, slower IAF and lower fronto-parietal connectivity in the lesioned hemisphere may underlie impaired perceptual processing in the contralateral hemispace.

### Alpha amplitude as a stable marker, while frequency and connectivity may reflect state-dependent processing

Nonetheless, in our case, these indices during rest did not consistently predict attention bias across different tasks. It could be that the alpha-amplitude changes versus alpha-frequency and -connectivity changes are more trait-like versus state-dependent. Specifically, while interhemispheric imbalance in alpha-amplitude could represent a stable marker of the attention bias, IAF and connectivity changes could mirror task-dependent stimulus-processing capacities. Indeed, this hypothesis has been recently hinted, as on-task alpha amplitude versus IAF and connectivity changes explained the behavior of the neglect patients through two separate model pathways, as identified via Structure Equation Model ^29^.

The first pathway linked pre-stimulus frontoparietal connectivity and individual alpha frequency to visual processing (N100) that In turn shaped perception, while the second one directly connected inter-hemispheric alpha-amplitude distribution to perceptual asymmetry. Therefore, it could be that measures of IAF and connectivity should be acquired during attention allocation to be maximally informative to task performance.

### Eyes-closed rs-EEG as more informative measure of baseline cortical activity compared to eyes-open condition

Most EEG differences between VSN patients and healthy controls emerged during the eyes-closed condition. In particular, alpha amplitude asymmetries observed exclusively during eyes-closed resting state consistently predicted attentional bias and perceptual deficits in VSN patients. These findings suggest that eyes-closed resting-state EEG may offer a more stable and reliable marker of attentional impairments in VSN than the eyes-open condition. This distinction likely reflects the different roles these conditions serve: eyes-closed EEG is better suited for assessing baseline arousal, as it captures intrinsic brain activity in the absence of visual input, whereas eyes-open EEG provides a baseline of activation, more relevant when comparing resting and task-evoked states^56^. Therefore, eyes-closed rs-EEG might yield more stable markers of physiological changes and symptom severity in VSN.

### Specificity of Alpha Effects Compared to Other Frequency Bands

Crucially, we also looked at other frequency bands, such as theta and beta, to affirm that the interhemispheric differences and their values as predictors of attention bias are specific for the alpha band. While the connectivity changes were not present across alternative frequencies, amplitude differences were present only in the beta-band. However, they did not consistently predict attention bias as seen for the alpha frequency. These control analyses A) argue against the notion that hemisphere differences result purely from structural brain damage, which would likely affect all frequencies, and (B) reinforce the dominant role of alpha oscillations in spatial attention.

To summarize, the current study provides a comprehensive overview of resting-state alpha oscillatory mechanisms in neglect patients and their role in attentional biases. First, we show that reduced alpha amplitude and frequency, along with diminished fronto-parietal connectivity in the right hemisphere, significantly diverge from the oscillatory patterns observed in healthy individuals—highlighting these alterations as potential electrophysiological markers of VSN. Second, we find that interhemispheric differences in alpha amplitude are a consistent marker of attention bias across different task contexts; supporting the notion that interhemispheric imbalance represents a core neural substrate of neglect.

Furthermore, our findings suggest that while alpha-amplitude asymmetry may serve as a stable biomarker of neglect-related deficits, alpha frequency and connectivity may be more reflective of state-dependent attentional processes. Overall, these findings contribute to the growing body of evidence supporting the role of interhemispheric interactions in spatial attention and may inform the development of targeted neurorehabilitation strategies. Future research should explore whether task-related measures of alpha frequency and connectivity could provide additional insights into attentional processing in neglect and whether interventions aimed at rebalancing alpha asymmetry could alleviate spatial attention deficits.

## Glossary

VSN: Visuospatial neglect
EEG: Electroencephalography
IAF: Individual Alpha Frequency
CVDT: Computerized visual detection task
SCT: Star Cancellation Task
SLBT: Schenkenberg Line Bisection Task

## Author Contributions

J. Trajkovic: drafting/revision of the manuscript for content; analysis or interpretation of data. M.M. Spanje: drafting/revision of the manuscript for content; major role in the acquisition of data. I. Leunissen: drafting/revision of the manuscript for content; analysis or interpretation of data. T. Schuhmann: drafting/revision of the manuscript for content; study concept or design; analysis or interpretation of data.

## References

1. Li K, Malhotra PA. Spatial neglect. Pract Neurol. 2015;15(5):333–339. doi:10.1136/practneurol-2015-001115

2. Corbetta M, Shulman GL. Spatial Neglect and Attention Networks. Annual Review of Neuroscience. 2011;34(Volume 34, 2011):569–599. doi:10.1146/annurev-neuro-061010-113731

3. Buxbaum LJ, Ferraro MK, Veramonti T, et al. Hemispatial neglect: Subtypes, neuroanatomy, and disability. Neurology. 2004;62(5):749–756. doi:10.1212/01.wnl.0000113730.73031.f4

4. Bartolomeo P. From competition to cooperation: Visual neglect across the hemispheres. Rev Neurol (Paris). 2021;177(9):1104–1111. doi:10.1016/j.neurol.2021.07.015

5. Bartolomeo P, Thiebaut de Schotten M, Chica AB. Brain networks of visuospatial attention and their disruption in visual neglect. Front Hum Neurosci. 2012;6:110. doi:10.3389/fnhum.2012.00110

6. Carter AR, Astafiev SV, Lang CE, et al. Resting Inter-hemispheric fMRI Connectivity Predicts Performance after Stroke. Ann Neurol. 2010;67(3):365–375. doi:10.1002/ana.21905

7. Stone SP, Halligan PW, Greenwood RJ. The incidence of neglect phenomena and related disorders in patients with an acute right or left hemisphere stroke. Age Ageing. 1993;22(1):46–52. doi:10.1093/ageing/22.1.46

8. Heilman KM, Abell TVD. Right hemisphere dominance for attention. Neurology. 1980;30(3):327–327. doi:10.1212/WNL.30.3.327

9. Kinsbourne M. Hemi-neglect and hemisphere rivalry. Adv Neurol. 1977;18:41–49.

10. Mesulam MM. A cortical network for directed attention and unilateral neglect. Ann Neurol. 1981;10(4):309–325. doi:10.1002/ana.410100402

11. Kastner S, Pinsk MA, De Weerd P, Desimone R, Ungerleider LG. Increased activity in human visual cortex during directed attention in the absence of visual stimulation. Neuron. 1999;22(4):751–761. doi:10.1016/s0896-6273(00)80734-5

12. Hopfinger JB, Buonocore MH, Mangun GR. The neural mechanisms of top-down attentional control. Nature Neuroscience. 2000;3(3):284–291. doi:10.1038/72999

13. Klimesch W, Sauseng P, Hanslmayr S. EEG alpha oscillations: the inhibition-timing hypothesis. Brain Res Rev. 2007;53(1):63–88. doi:10.1016/j.brainresrev.2006.06.003

14. Klimesch W, Doppelmayr M, Russegger H, Pachinger T, Schwaiger J. Induced alpha band power changes in the human EEG and attention. Neurosci Lett. 1998;244(2):73–76. doi:10.1016/s0304-3940(98)00122-0

15. Trajkovic J, Gregorio FD, Avenanti A, Thut G, Romei V. Two oscillatory correlates of attention control in the alpha-band with distinct consequences on perceptual gain and metacognition. J Neurosci. Published online April 5, 2023. doi:10.1523/JNEUROSCI.1827-22.2023

16. Thut G, Nietzel A, Brandt SA, Pascual-Leone A. α-Band Electroencephalographic Activity over Occipital Cortex Indexes Visuospatial Attention Bias and Predicts Visual Target Detection. J Neurosci. 2006;26(37):9494–9502. doi:10.1523/JNEUROSCI.0875-06.2006

17. Bartolomeo P, Thiebaut de Schotten M, Doricchi F. Left unilateral neglect as a disconnection syndrome. Cereb Cortex. 2007;17(11):2479–2490. doi:10.1093/cercor/bhl181

18. Romei V, Brodbeck V, Michel C, Amedi A, Pascual-Leone A, Thut G. Spontaneous fluctuations in posterior alpha-band EEG activity reflect variability in excitability of human visual areas. Cereb Cortex. 2008;18(9):2010–2018. doi:10.1093/cercor/bhm229

19. Di Gregorio F, Trajkovic J, Roperti C, et al. Tuning alpha rhythms to shape conscious visual perception. Current Biology. 2022;32(5):988–998.e6. doi:10.1016/j.cub.2022.01.003

20. Coldea A, Veniero D, Morand S, et al. Effects of Rhythmic Transcranial Magnetic Stimulation in the Alpha-Band on Visual Perception Depend on Deviation From Alpha-Peak Frequency: Faster Relative Transcranial Magnetic Stimulation Alpha-Pace Improves Performance. Front Neurosci. 2022;16:886342. doi:10.3389/fnins.2022.886342

21. Samaha J, Postle BR. The Speed of Alpha-Band Oscillations Predicts the Temporal Resolution of Visual Perception. Current Biology. 2015;25(22):2985–2990. doi:10.1016/j.cub.2015.10.007

22. Bertaccini R, Ellena G, Macedo-Pascual J, et al. Parietal Alpha Oscillatory Peak Frequency Mediates the Effect of Practice on Visuospatial Working Memory Performance. Vision (Basel). 2022;6(2):30. doi:10.3390/vision6020030

23. Trajkovic J, Di Gregorio F, Thut G, Romei V. Transcranial magnetic stimulation effects support an oscillatory model of ERP genesis. Current Biology. 2024;34(5):1048–1058.e4. doi:10.1016/j.cub.2024.01.069

24. Cecere R, Rees G, Romei V. Individual differences in alpha frequency drive crossmodal illusory perception. Curr Biol. 2015;25(2):231–235. doi:10.1016/j.cub.2014.11.034

25. Cooke J, Poch C, Gillmeister H, Costantini M, Romei V. Oscillatory Properties of Functional Connections Between Sensory Areas Mediate Cross-Modal Illusory Perception. J Neurosci. 2019;39(29):5711–5718. doi:10.1523/JNEUROSCI.3184-18.2019

26. Sauseng P, Klimesch W, Stadler W, et al. A shift of visual spatial attention is selectively associated with human EEG alpha activity. European Journal of Neuroscience. 2005;22(11):2917–2926. doi:10.1111/j.1460-9568.2005.04482.x

27. Ebisu T, Fukunaga M, Murase T, et al. Functional Connectivity Pattern Using Resting-state fMRI as an Assessment Tool for Spatial Neglect during the Recovery Stage of Stroke: A Pilot Study. Magn Reson Med Sci. 2023;22(3):313–324. doi:10.2463/mrms.mp.2022-0010

28. Dubovik S, Pignat JM, Ptak R, et al. The behavioral significance of coherent resting-state oscillations after stroke. NeuroImage. 2012;61(1):249–257. doi:10.1016/j.neuroimage.2012.03.024

29. Di Gregorio F, Petrone V, Casanova E, et al. Hierarchical psychophysiological pathways subtend perceptual asymmetries in Neglect. Neuroimage. 2023;270:119942. doi:10.1016/j.neuroimage.2023.119942

30. Puig-Pijoan A, Giralt-Steinhauer E, Zabalza de Torres A, et al. Underdiagnosis of Unilateral Spatial Neglect in stroke unit. Acta Neurol Scand. 2018;138(5):441–446. doi:10.1111/ane.12998

31. Azouvi P, Bartolomeo P, Beis JM, Perennou D, Pradat-Diehl P, Rousseaux M. A battery of tests for the quantitative assessment of unilateral neglect. Restor Neurol Neurosci. 2006;24(4-6):273–285.

32. Rihs TA, Michel CM, Thut G. Mechanisms of selective inhibition in visual spatial attention are indexed by alpha-band EEG synchronization. Eur J Neurosci. 2007;25(2):603–610. doi:10.1111/j.1460-9568.2007.05278.x

33. Rihs TA, Michel CM, Thut G. A bias for posterior alpha-band power suppression versus enhancement during shifting versus maintenance of spatial attention. Neuroimage. 2009;44(1):190–199. doi:10.1016/j.neuroimage.2008.08.022

34. Middag-van Spanje M, Schuhmann T, Nijboer T, van der Werf O, Sack AT, van Heugten C. Study protocol of transcranial electrical stimulation at alpha frequency applied during rehabilitation: A randomized controlled trial in chronic stroke patients with visuospatial neglect. BMC Neurology. 2022;22(1):402. doi:10.1186/s12883-022-02932-7

35. Schuhmann T, Duecker F, Middag-van Spanje M, et al. Transcranial alternating brain stimulation at alpha frequency reduces hemispatial neglect symptoms in stroke patients. Int J Clin Health Psychol. 2022;22(3):100326. doi:10.1016/j.ijchp.2022.100326

36. Getzmann S, Gajewski PD, Schneider D, Wascher E. Resting-state EEG data before and after cognitive activity across the adult lifespan and a 5-year follow-up. Sci Data. 2024;11(1):988. doi:10.1038/s41597-024-03797-w

37. Duecker F, Schuhmann T, Bien N, Jacobs C, Sack AT. Moving Beyond Attentional Biases: Shifting the Interhemispheric Balance between Left and Right Posterior Parietal Cortex Modulates Attentional Control Processes. J Cogn Neurosci. 2017;29(7):1267–1278. doi:10.1162/jocn_a_01119

38. Schuhmann T, Kemmerer SK, Duecker F, et al. Left parietal tACS at alpha frequency induces a shift of visuospatial attention. PLoS One. 2019;14(11):e0217729. doi:10.1371/journal.pone.0217729

39. Watson AB, Pelli DG. Quest: A Bayesian adaptive psychometric method. Perception & Psychophysics. 1983;33(2):113–120. doi:10.3758/BF03202828

40. Wilson B, Cockburn J, Halligan P. Development of a behavioral test of visuospatial neglect. Arch Phys Med Rehabil. 1987;68(2):98–102.

41. Dalmaijer ES, Van der Stigchel S, Nijboer TCW, Cornelissen THW, Husain M. CancellationTools: All-in-one software for administration and analysis of cancellation tasks. Behav Res. 2015;47(4):1065–1075. doi:10.3758/s13428-014-0522-7

42. Trajkovic J, Sack AT, Romei V. EEG-based biomarkers predict individual differences in TMS-induced entrainment of intrinsic brain rhythms. Brain Stimul. 2024;17(2):224–232. doi:10.1016/j.brs.2024.02.016

43. Cohen M. Effects of time lag and frequency matching on phase-based connectivity. J Neurosci Methods. 2015;250:137–146. doi:10.1016/j.jneumeth.2014.09.005

44. Trajkovic J, Di Gregorio F, Ferri F, Marzi C, Diciotti S, Romei V. Resting state alpha oscillatory activity is a valid and reliable marker of schizotypy. Sci Rep. 2021;11:10379. doi:10.1038/s41598-021-89690-7

45. Çiçek M, Nalçacı E. Interhemispheric asymmetry of EEG alpha activity at rest and during the Wisconsin Card Sorting Test: relations with performance. Biological Psychology. 2001;58(1):75–88. doi:10.1016/S0301-0511(01)00103-X

46. Snyder DB, Schmit BD, Hyngstrom AS, Beardsley SA. Electroencephalography resting-state networks in people with Stroke. Brain and Behavior. 2021;11(5):e02097. doi:10.1002/brb3.2097

47. Marcu GM, Szekely-Copîndean RD, Radu AM, et al. Resting-state frontal, frontlateral, and parietal alpha asymmetry:A pilot study examining relations with depressive disorder type and severity. Front Psychol. 2023;14. doi:10.3389/fpsyg.2023.1087081

48. Pietrelli M, Zanon M, Làdavas E, Grasso PA, Romei V, Bertini C. Posterior brain lesions selectively alter alpha oscillatory activity and predict visual performance in hemianopic patients. Cortex. 2019;121:347–361. doi:10.1016/j.cortex.2019.09.008

49. Zhang JJ, Bai Z, Fong KNK. Resting-state cortical electroencephalogram rhythms and network in patients after chronic stroke. J NeuroEngineering Rehabil. 2024;21(1):32. doi:10.1186/s12984-024-01328-7

50. Baldassarre A, Ramsey L, Hacker CL, et al. Large-scale changes in network interactions as a physiological signature of spatial neglect. Brain. 2014;137(Pt 12):3267–3283. doi:10.1093/brain/awu297

51. Doesburg SM, Green JJ, McDonald JJ, Ward LM. From local inhibition to long-range integration: A functional dissociation of alpha-band synchronization across cortical scales in visuospatial attention. Brain Research. 2009;1303:97–110. doi:10.1016/j.brainres.2009.09.069

52. Hembrook Short JR, Mock VL, Briggs F. Attentional modulation of neuronal activity depends on neuronal feature selectivity. Curr Biol. 2017;27(13):1878–1887.e5. doi:10.1016/j.cub.2017.05.080

53. Hembrook-Short JR, Mock VL, Usrey WM, Briggs F. Attention Enhances the Efficacy of Communication in V1 Local Circuits. J Neurosci. 2019;39(6):1066–1076. doi:10.1523/JNEUROSCI.2164-18.2018

54. Luzio PD, Tarasi L, Silvanto J, Avenanti A, Romei V. Human perceptual and metacognitive decision-making rely on distinct brain networks. PLOS Biology. 2022;20(8):e3001750. doi:10.1371/journal.pbio.3001750

55. Chiappini E, Silvanto J, Hibbard PB, Avenanti A, Romei V. Strengthening functionally specific neural pathways with transcranial brain stimulation. Curr Biol. 2018;28(13):R735–R736. doi:10.1016/j.cub.2018.05.083

56. Barry RJ, De Blasio FM. EEG differences between eyes-closed and eyes-open resting remain in healthy ageing. Biological Psychology. 2017;129:293–304. doi:10.1016/j.biopsycho.2017.09.010

